# Comparison of the Gut Microbiota Composition Between Captive and Wild Roe Deer

**DOI:** 10.1101/831222

**Authors:** Jinyue Liu, Xue Liang, Yanhua Liu

## Abstract

In this paper, 16S-rRNA gene Illumina HiSeq sequencing was used to analyze the structural diversity of captive and wild roe deer gut flora. The results show that the microbial diversity in the feces of wild roe deer is higher than in that of captive roe deer. Both roe deer have similar flora at the phylum level, but the main genus has significant differences. The microbial group that plays an important role in captive roe deer is Bacteroidetes; in wild roe deer it is Firmicutes. This difference is mainly due to the differences in living environment, diet, and physiological functions of the two groups. In conclusion, our study makes people have a better understanding of the intestinal flora of roe deer. By comparing the intestinal microbial structure differences between captive and wild roe deer, it provides theoretical basis for people to raise captive roe deer and provides reference for the protection of wild roe deer.

**IMPORTANCE:** Many studies have shown that large and complex microbes in the gut of humans and non-human animals, intestinal microbes are thought to co-evolve with the host, help the host acquire nutrients, regulate immunity and to help maintain host homeostasis. The roe deer (*Capreolus spp.*) is a ruminant. Wild roe deer are listed on the *List of Terrestrial Wild Animals Protected by the State or Have Important Economic and Scientific Values*, wild roe deer is also a Chinese national protected animal under second class protection. However, current research on the gut microbiota of roe deer has not been reported.

---

The microbiota in the intestine is not fixed, and changes with the age, diet, and living environment of the host (1). Numerous studies have shown that the occurrence of intestinal diseases in mammals is related to intestinal microbes, which have a great influence on the normal physiological activities of the host(2). Especially for ruminants, due to their unique digestive characteristics, the microbiota is good for digesting foods with high fiber content, but at the same time it is more susceptible to many diseases(3). Therefore, the microflora in the intestinal tract is more important and meaningful for ruminants.

The roe deer (*Capreolus spp.*) is a ruminant belonging to the Cervidae family. There are two species of roe deer: the Siberian roe deer (*Capreolus pygargus*) and the European roe deer (*Capreolus capreolus*)(4).*C. pygargus* is widely distributed in Northeast Asia, including Russia, Central Asia, Korean Peninsula and China, especially the northeast of China. The roe deer not only has edible value, but also has medicinal value and ornamental value(5). There is a legend: If you eat meat, you will become a fairy. For this reason, people have excessive poaching, and the wild resources of the roe deer are scarce. The roe deer are the main prey of large cats such as tigers and leopards(6–8). They play an important role in affecting their density and distribution. They also play an important role in maintaining ecosystem balance. So wild roe deer are listed in the *List of Terrestrial Wild Animals Protected by the State or Have Important Economic and Scientific Values*.

Some scholars have done a lot of research on roe deer. Some people have studied the hunting behavior of roe deer(9), some people have studied the diet of wild roe deer(10), and some experts have analyzed the immunohistochemistry of roe deer blood cells(11). However, studies on gut microbes have not been reported. This paper compares the composition and structure of microbes in the feces of different genders and different growth environments, and contributes to the study of intestinal microbes, providing a scientific basis for the cultivation of captive roe deer. At the same time, it provides ideas for protecting wild roe deer.

## RESULTS

### Analysis of captive and wild roe deer microbiota based on 16SrRNA sequencing results

We divided into two groups, A and B, according to the two living conditions. The number of shared OTUs in the two groups was 2717, and the number of shared OTUs in the captive (A) and wild (B) groups was 34,295 respectively(Figure 1)

**FIGURE 1.**
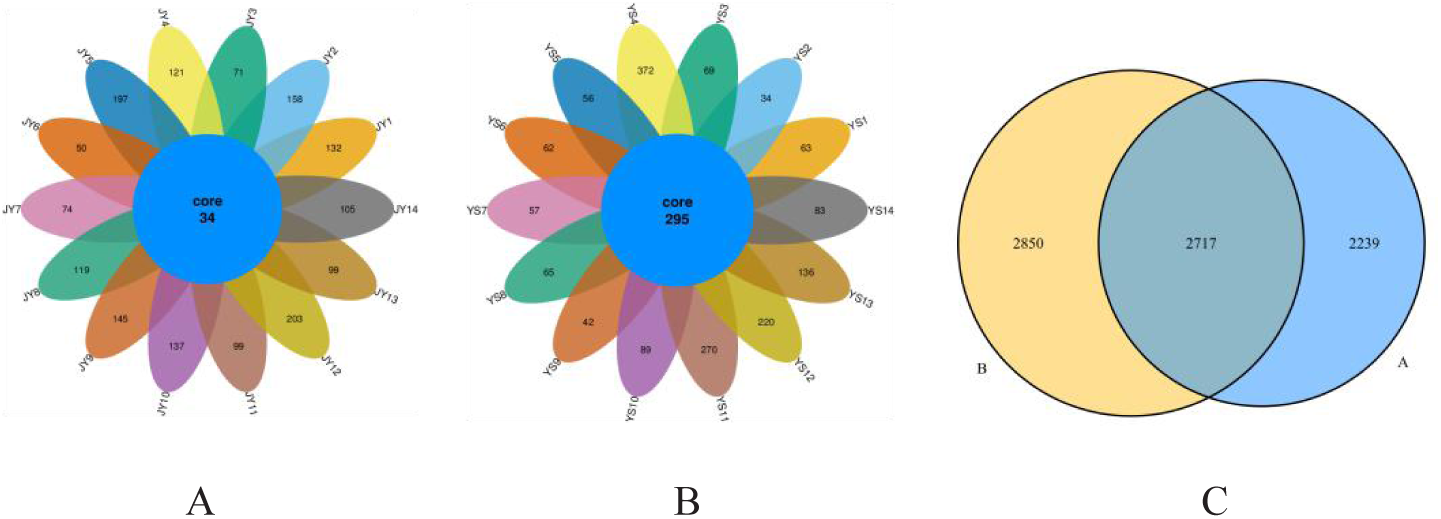
Venn diagrams. The Venn diagrams show the numbers of OTUs (97% sequence identity) that were shared or not shared by captive and wild individuals, respectively, depending of overlaps. (A) The number of OTUs shared by captive. (B) The number of OTUs shared by wild. (C) The number of OTUs shared by captive and wild.

In order to investigate the species composition of the sample and predict the species abundance in the sample, we produced species accumulation curve to determine if the sample size is sufficient and to estimate the species richness (Figure 2). When the sample size is 0-10, as the sample size increases, the curve rises sharply, indicating that a large number of species are found in the community. When the sample size is about 25, the curve tends to be flat. It is indicated that the microbial species in the intestine do not increase significantly with the increase in the number of fecal samples beyond this point. It is therefore, suggested that the experiment was adequately sampled and representative, and further data analysis can justifiably be performed.

**FIGURE 2.**
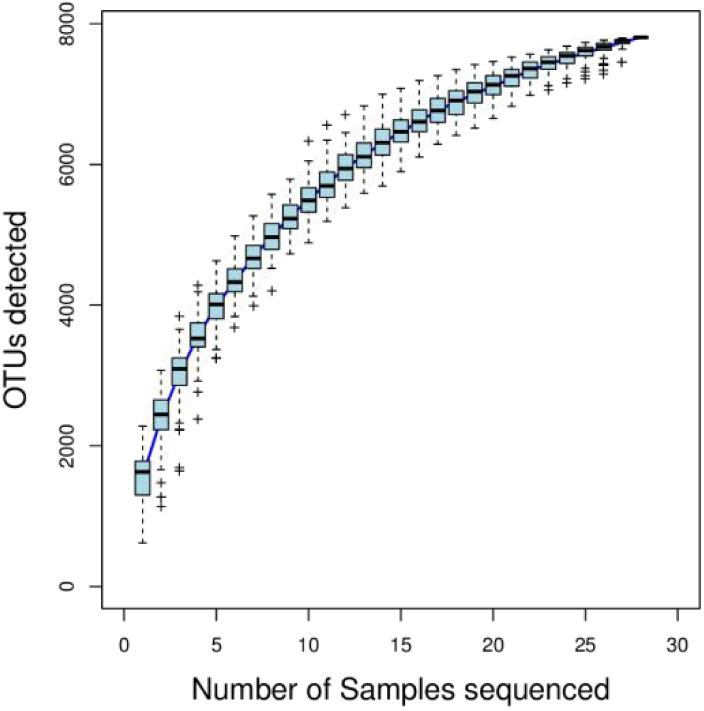
Species accumulation curve. The x-axis shows the number of samples and the y-axis shows the OTUs after sampling(operational taxonomic units, OTUs). A sharp rise in the curve indicates that the sample amount is insufficient, and the sampling amount needs to be increased. When the curve tends to be gentle, it indicates that the sampling is sufficient and data analysis can be performed.

This study also used the Rarefaction Curve to describe the sample diversity within the group (Figure 3). The results showed that as the depth of sequencing increased, the number of observed species increased, and the end of the curve gradually became flat with the increase of the number of sequences in the sample.

**FIGURE 3.**
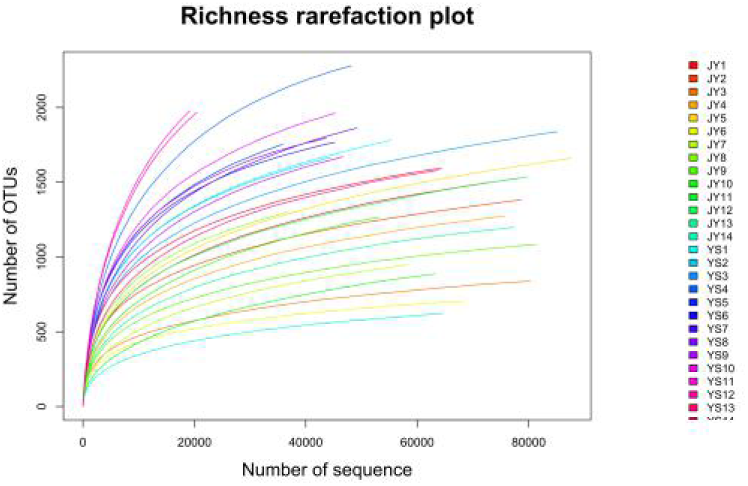
Rarefaction curves. The x-axis shows the number of valid sequences per sample and the y-axis shows the observed species(operational taxonomic units, OTUs). Each curve in the graph represents a different sample and is shown in a different color. As the sequencing depth increased, the number of OTUs also increased. Eventually the curves began to plateau, indicating that as the number of extracted sequences increased,the number of OTUs detected was decreased.

### Differences in Alpha-diversity between captive and wild roe deer gut microbes

The diversity of captive and wild roe deer gut microbes was compared by Alpha-diversity index analyses (Shannon, ACE, Chao1, coverage, Simpson) and beta-diversity analyses (Unweighted and Weighted Unifrac distance matrix). The results of Alpha-diversity analysis showed that the Shannon values of captive and wild roe deer were 4.85±0.49 and 5.63±0.20 (P<0.01), respectively, ACE were 1501.28±417.61 and 2254.00±253.82 (P<0.05), respectively, Chao1 were 1486.15±425.48and 2245.87±216.34 (P<0.01), respectively, coverage was 1.00±0.00 and 1.00±0.01 (P<0.05), respectively and for Simpson all were 0.02±0.01 (P>0.05)(Table 2). The coverage value indicates that the data obtained in this study is a good response to the bacterial diversity of the sample.

### Differences in Beta-diversity between captive and wild roe deer gut microbes

The Beta-diversity results showed that there were also differences in intestinal microbes between the A and B groups. Unweighted and Weighted Unifrac distance drawn Heatmap shows the difference between the two samples(Figure 4). Through the analysis of non-metric multi-dimensional scaling (NMDS) based on OTU level(Figure 5), there was significant separation between the two groups, the A group was relatively dispersed, the difference between the samples was large, and the B group was relatively concentrated.

**FIGURE 4.**
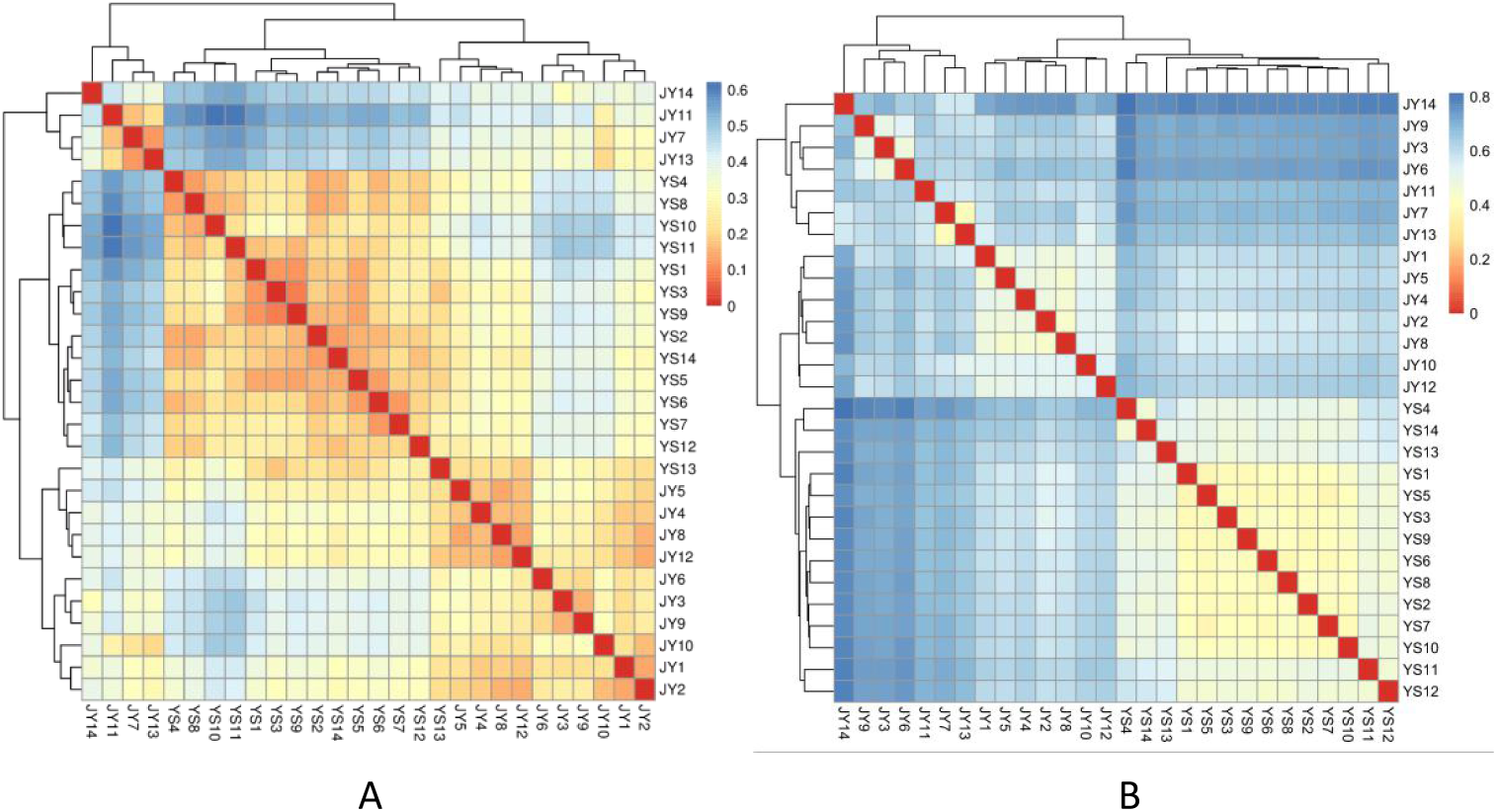
Heatmap of β diversity index between the two groups. (A) Weighted Unifrac distance drawn Heatmap. (B) Unweighted Unifrac distance drawn Heatmap. The color block represents the distance value. The redder the color is, the closer the samples are, the higher the similarity is, and the bluer the distance is. In the heat map, the distance between samples can be seen through the clustering tree.

**FIGURE 5.**
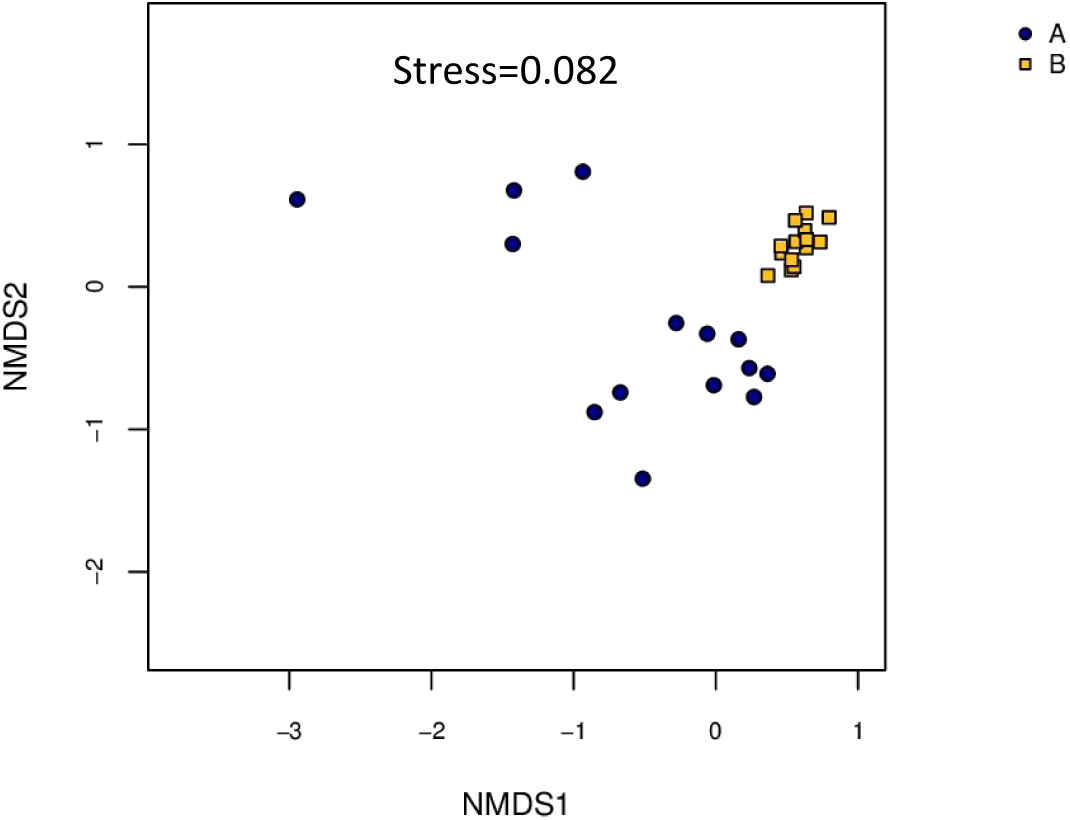
Non-metric multidimensional scaling (NMDS) analysis. Each point in the graph represents one sample, and different colors represent different groups. The distance between points represents the level of difference. Stress lower than 0.2 indicates that the NMDS analysis is reliable. The closer the samples in the graph, the higher their similarity.

Anosim analysis demonstrated differences in intestinal microbes between captive and wild roe deer (R = 0.385, P = 0.001)(Figure 6). The intra-group difference between captive and wild roe deer microbes was less than the difference between groups, and the composition of intestinal microbes in the two groups was significantly different (P<0.05). The results of principal component analysis based on OTU levels in group A and group B gut intestinal flora show that both groups of roe deer have their own core flora. The contribution rates of the first principal component and the second principal component based on the Principal Component Analysis(PCA) were 36% and 12%(Figure 7), respectively. Wild and captive roe deer are apparently clustered together in their respective groups. PCoA analysis(Figure 8) based on Weighted Unifrac and Unweighted Unifrac distances clearly shows that there is an obvious analysis phenomenon in the two groups of samples, but the structural similarity of the wild roe deer group is relatively high.

**FIGURE 6.**
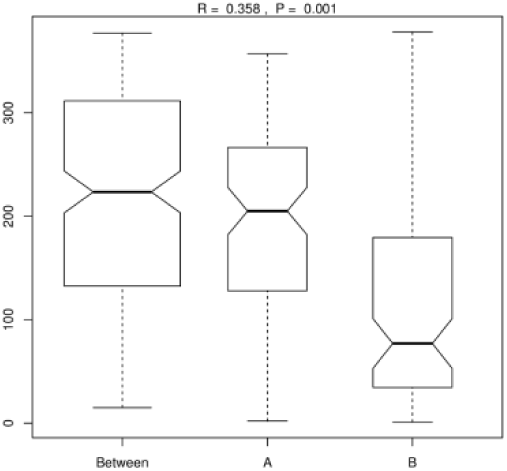
ANOSIM analysis. R-value: R-value range (–1, 1). Actual results are generally between 0 and 1. An R-value close to 0 represents no significant inter-group and intra-group differences. A R-value close to 1 shows that inter-group differences are greater than intra-group differences. P-value: The P-value represents the confidence level of the statistical analysis; P < 0.05 reflects a statistically significant difference. The y-axis represents the distance rank between samples, and the x-axis represents the results between both groups. Intra-group results are shown for each group. In the plot, the R-value was close to 1, indicating that inter-group differences were greater than the intra-group differences, and P < 0.05 shows that this result was statistically significant.

**FIGURE 7.**
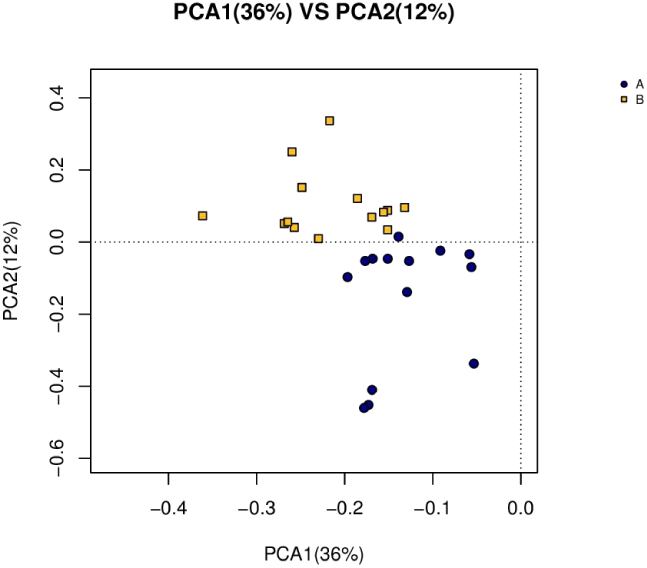
Principal Component Analysis diagram based on OTU. The different colors in the figure represent the samples in different groups, and the higher the similarity between the samples, the larger the aggregation in the figure.

**FIGURE 8.**
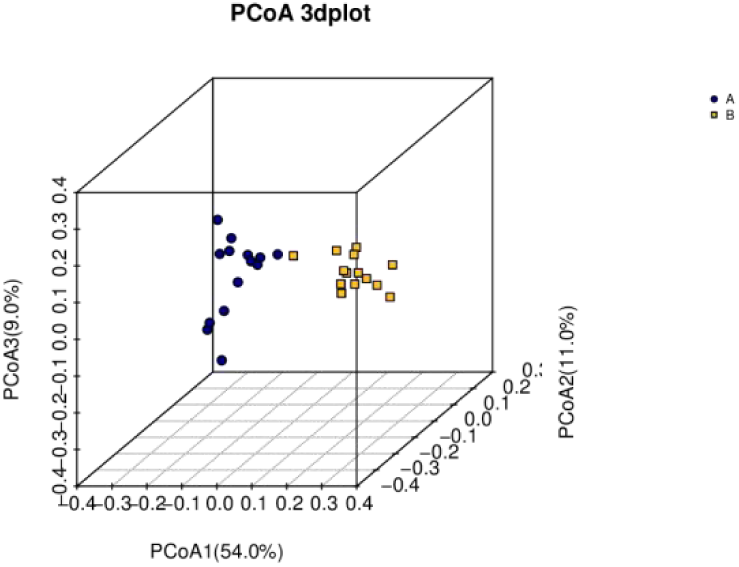
3D Principal Co-ordinates Analysis diagram based on unifrac distance matrix. Points of different colors represent samples in different groups. The higher the similarity between samples is, the larger the aggregation in the figure will be; on the contrary, the lower the similarity between samples is, the farther the spatial distance will be.

### Species analysis of differences in intestinal flora between captive and wild roe deer

Through the T-test, the species with significant differences (p < 0.05) at the phylum level were found. Within 95% confidence interval, the most significant differences were Candidatus Saccharibacteria, Lentisphaerae, Firmicutes, Actinobacteria, and Bacteroidetes. Through the T-test, the genus with significant difference in the abundance of the wild roe deer and the captive roe deer was found. The intestinal flora of captive roe deer is only significantly higher in the abundance of Clostridium XlVa than in the wild group (Figure 9)

To analyze whether there was a statistically significant difference between the wild and captive roe deer, LEfSe was used to establish species with significant differences between the groups. From the histogram of LDA value distribution, Groups A and B have populations with significantly different abundances. There were 40 species with abundance difference in group A and 35 species with abundance difference in group B. From the evolutionary branch map (Figure 10), it is clear that the microbial group that plays an important role in group A is Bacteroidetes. "radiation" inward to Bacteroidia, Bacteroidales. The microbial group that plays an important role in group B is Firmicutes, as well as Clostridia and Clostridiales.

**FIGURE 9.**
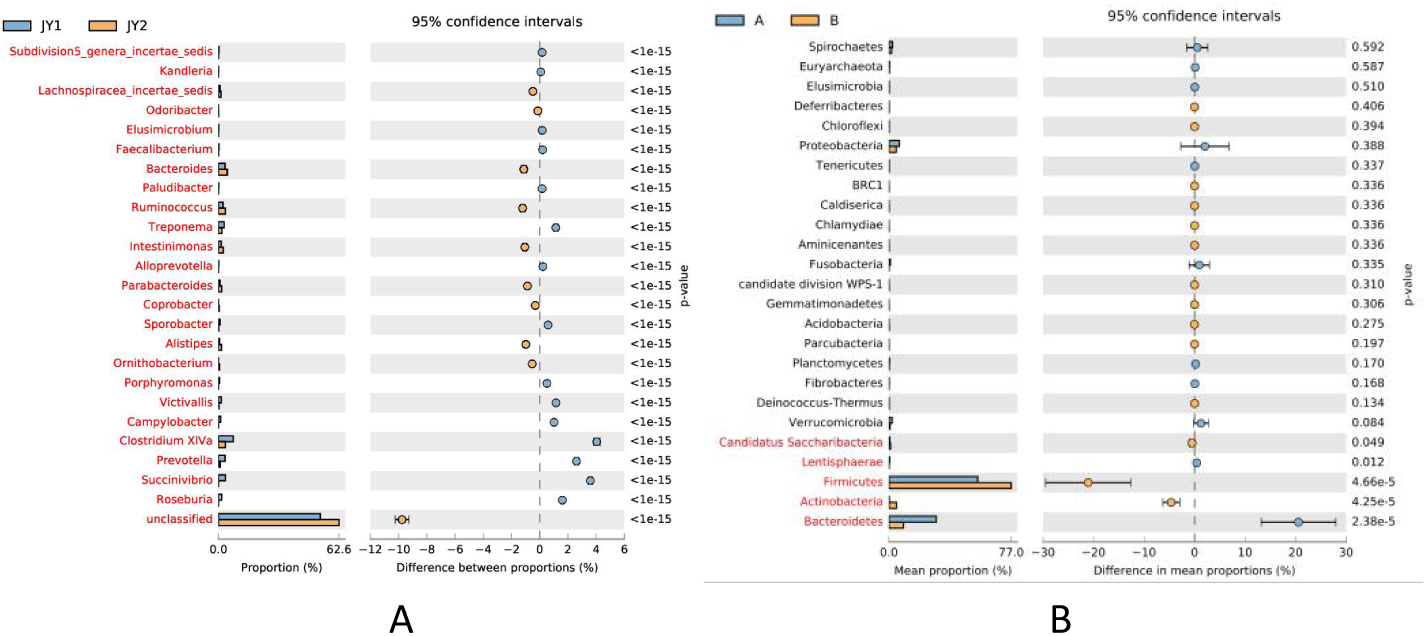
Difference comparison error diagram. (A) Analysis of species differences at genus level between groups based on T-test. (B) Analysis of species differences at phylum level between groups based on T-test. The left figure shows the abundance proportion of different species classification in two samples (groups). The middle figure shows the difference proportion of species classification abundance within the 95% confidence interval. The rightmost value is p value, p value < 0.05 indicates significant difference. (only list the 25 lowest p values)

**FIGURE 10.**
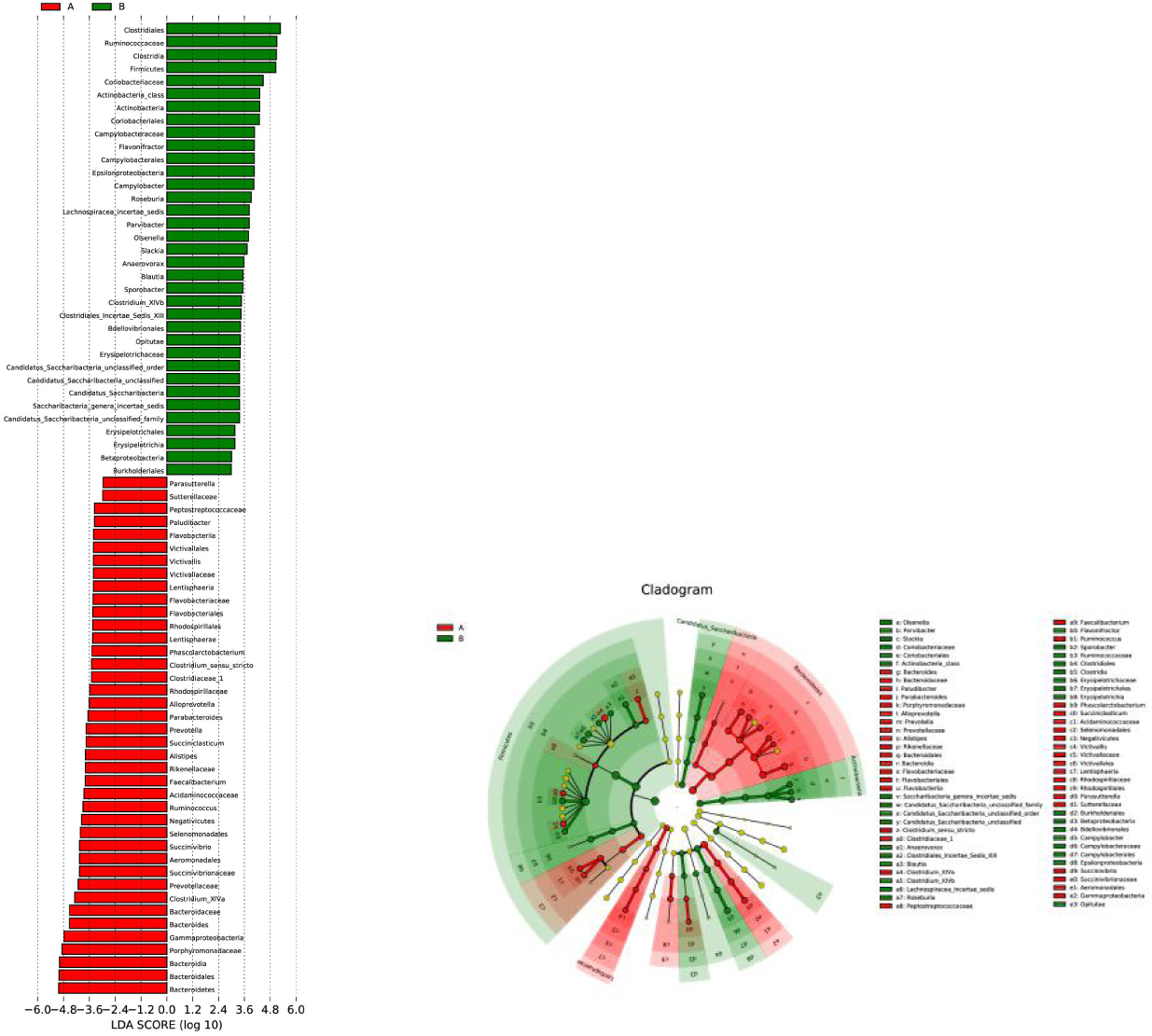
LEfSe analysis. (A) Plot from LEfSe analysis. The plot was generated using the online LEfSe project. The length of the bar column represents the LDA score. The figure shows the microbial taxa with significant differences between the captive (red) and wild (green) (LDA score > 2). (B) A cladogram showing the differences in relative abundance of taxa at five levels between captive and wild. The plot was generated using the online LEfSe project. The red and green circles mean that captive and wild showed differences in relative abundance and yellow circles mean non-significant differences.

## DISCUSSION

This experiment used 16S-rRNA gene Illumina HiSeq sequencing analysis. The composition and structure of captive and wild roe deer gut microbes were studied. The results show that there are differences in both the composition and structure of microbial flora in captive and wild roe deer intestines. The core microbial flora of captive and wild roe deer were found to be mainly Bacteroidetes and Firmicutes. This result is consistent with the results of some of the ruminant gut microbe studied by earlier works. Sundset *et al.* show 76.7% of the intestinal flora of Svalbard reindeer (*Rangifer tarandus platyrhynchus*) are Firmicutes and 22.5% are Bacteriodales (12). The results of Gruninger *et al.* show that Bacteroidetes and Firmicutes are the main intestinal microflora(13). The results of Ishaq *et al.* also show that Bacteroidetes and Firmicutes are dominant in the intestinal flora of many ruminants(14).

The digestive capacity of human and animal individuals often depends on the composition of their gut microflora(15–18). When the relative abundance of Firmicutes is high, the ability of mice to harvest energy from food becomes stronger(17). When Bacteroidetes have a higher abundance, the ability of mice to obtain carbohydrates or fats from the diet is limited(19). Some scholars have confirmed that the proportion of Bacteroidetes and Firmicutes in the intestine is related to fat deposition(20). Administration of the Anti-obesity complex for 8 weeks into obese mice, Bacteroidetes increase in relative abundance, the relative abundance of Firmicuts is reduced(21), it is shown that intestinal microbes with a low B/F ratio have higher fermentation efficiency and more energy from food, thus promoting fat deposition. In the wild, the roe deer is cold in winter and lacks food. It consumes a lot of energy to forage and escape natural predators, as well as maintaining body temperature. The structure of the high Firmimuts in the intestines helps them absorb energy from food as much as possible to maintain their body’s needs. Studies have shown that long-term diet is closely related to intestinal microbial components, and people with more protein and fat in the diet have more Bacteroidetes in the intestine(22). The food in the roe deer diet is mainly a variety of grains and grasses. It is rarely active every day, and does not consume a lot of energy. Therefore, Bacteroidetes is more abundant in the captive roe deer. In summary, diet affects the gut microbes of the roe deer to a certain extent. For captive roe deer, manufactured animalfeed is their main source of food. The food they need throughout their life comes from human supply, so their growth, reproduction, and health are all related to diet. For the wild roe deer, their intestinal microbes are formed through centuries of evolution and are relatively stable(1).

From the perspective of alpha-diversity analysis,the values of shannon and Chao1 can be seen by T test, the difference is extremely significant, ACE are significantly different, and simpson has no significant difference,the diversity of microbial communities in the wild gut is higher than that in captivity, This is consistent with the structure studied by Hu *et al.*(3). Many scholars believe that wild species contain comparatively more complex and richer intestinal flora(23–26). The captive roe deer has a relatively stable living environment. The food is fixed for various grains and hay materials, while the wild roe deer food is complex and diverse, and the amount of exercise and energy consumption is large. Therefore, the microbial diversity of the wild roe deer is higher. From the perspective of beta-diversity analysis, PCA, PCOA, and NMDS all showed significant separation between captive and wild roe deer fecal samples. This is related to the diet, digestion, physiological functions, lifestyle, etc.(2). The families of the bacteria with significant differences were found by LEfSe analysis and T-test. The microbial group that plays an important role in captivity is Bacteroidia and Bacteroidales of Bacteroidetes. The microbial group that plays an important role in the wild is Clostridia and Clostridiales of Firmicutes. Bacteroidales is one of the important dominant strains in the gut of mammals and has the function of helping to digest carbohydrates. Clostridiales is one of the dominant strains in gut microbes and contains multiple strains of fermented sugar, polyhydric alcohols, amino acid, organic acid, 1H-Purine and other organic compounds.

In addition to Bacteroidetes and Firmicuts, the most common bacteria in the gut microbiota are Candidatus Saccharibacteria, Lentisphaerae and Actinobacteria. Candidatus Saccharibacteria has the obligate fermentation metabolism, fermentation glucose and other sugars, and produce lactic acid; it belongs to Gram-positive (27). Lentisphaerae is Gram-negative, non-motile, non-pigmented and strictly aerobic(28). Actinomycete is one of the oldest bacterial phylum and plays an important role in medicine and biotechnology (29); it belongs to Gram-positive. In addition to regulating the nutrition and metabolism of the host, intestinal microbes are also critical to the function and development of the immune system(30). Previous studies found that symbiotic bacteria present in the gut are sufficient to induce low-grade inflammation in neonates and sterile mice(31,32). Gram-positive bacteria, which produce exotoxin, cause disease(33), while Gram-negative bacteria, which produce endotoxin, cause disease by endotoxin(34). The reduction of Gram-positive bacteria and the increased oxygen tolerance of Gram-negative bacteria suggest an important role for the innate immune system in the formation of human intestinal microbes.

At the level of genus, it was observed that 43.2% of the sequences in captivity could not be classified into any known genus, and 60.5% of the sequences in the wild, indicating that it is likely to be a new bacterium, presently undescribed. This result is similar to the results of the gut microbiological studies of wild(13) and captive(35) ruminants. It proves that there are more flora in the intestines than in the past. Unclassified sequences and their phylogenetic conditions will be one of the important studies in the future(Hu et al.,2017).

## MATERIALS AND METHODS

### Sample Collection

The experiment collected 28 fecal samples from 28 roe deer(Table 1), 14 of which were captive, taken from Longjia Village, Shangying Town, Shulan City, Jilin City, Jilin Province. The feed for captive roe deer is mainly composed of corn, soybean and vanilla. All captive roe deer have been healthy for the previous three months without any antibiotics or anti-inflammatory drugs. The animal housing was thoroughly cleaned the night before collecting the feces. Roe deer usually produce feces in the morning. Fresh excreted feces was collected. The feces was obviously granular and the color was black or dark green. The samples were collected with a disposable sterile glove to avoid human contamination, and the samples were immediately poured into a sterile centrifuge tube with 95 % alcohol and sealed to avoid cross-contamination between the samples. The other 14 roe deer were wild. They were taken from Muling City, Heilongjiang Province. We followed the footprints of the wild roe deer and carried out strict distance separation, to ensure that the collection of each fecal sample likely belonged to different individuals. Feces was kept fresh and clean as much as possible, storing at −20 °C. Immediately after taking it back, it was put into the refrigerator for the next experimental processes.

**TABLE 1.**
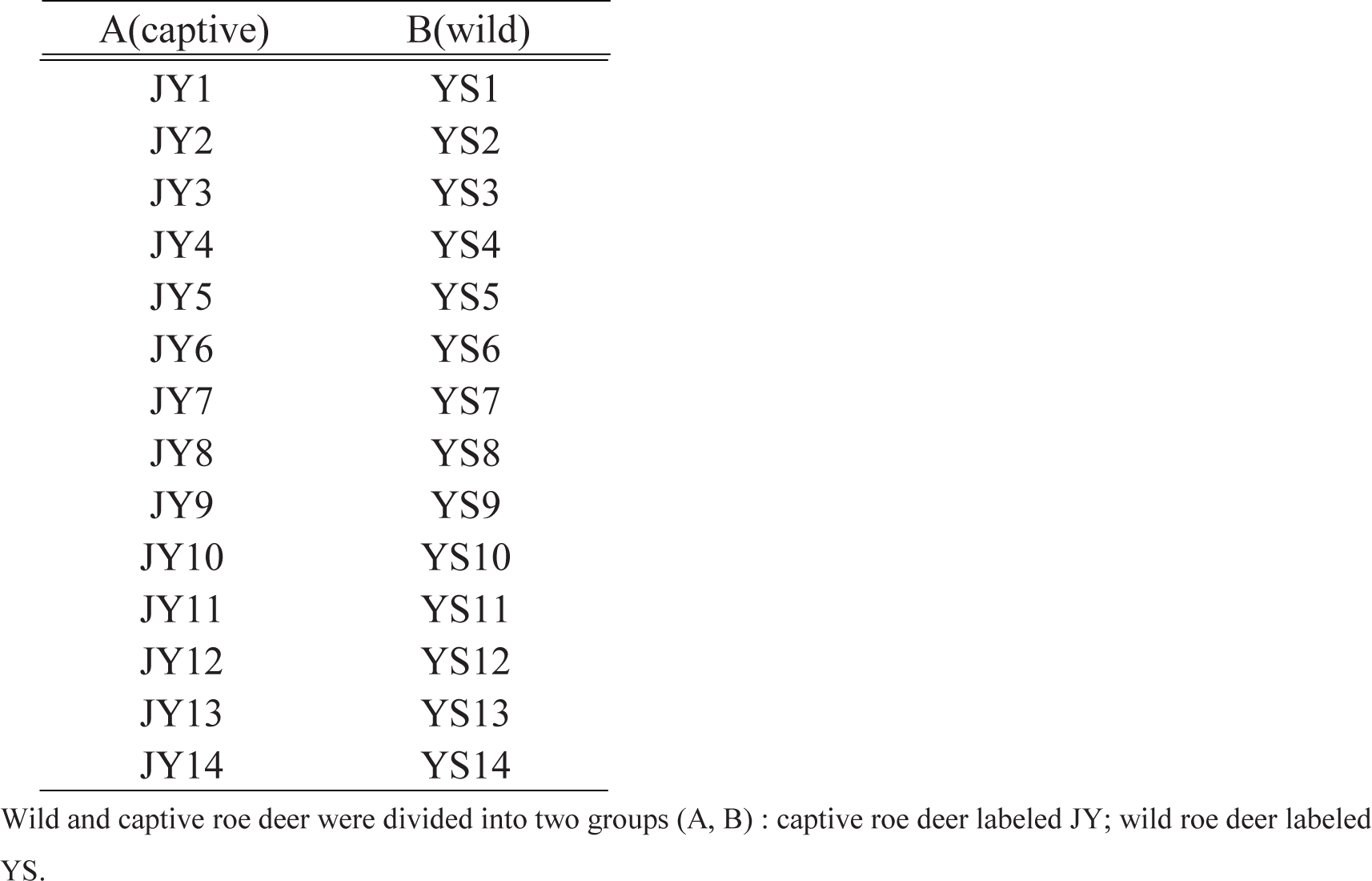
Group information on captive and wild roe deer

**TABLE 2.** Diversity index table for each sample

| Sample_ID | OTU_num | Shannon | ACE | Chao1 | Coverage | Simpson |
| --- | --- | --- | --- | --- | --- | --- |
| JY1 | 1528 | 5.24 | 1990.85 | 2007.20 | 0.99 | 0.02 |
| JY10 | 1266 | 5.19 | 1703.43 | 1618.74 | 0.99 | 0.01 |
| JY11 | 886 | 4.10 | 1277.39 | 1221.11 | 1.00 | 0.04 |
| JY12 | 1536 | 5.40 | 1999.60 | 2007.48 | 0.99 | 0.01 |
| JY13 | 1198 | 4.79 | 1531.33 | 1553.96 | 1.00 | 0.03 |
| JY14 | 620 | 3.79 | 772.98 | 785.28 | 1.00 | 0.06 |
| JY2 | 1382 | 5.23 | 1798.16 | 1847.71 | 1.00 | 0.02 |
| JY3 | 838 | 4.73 | 1074.70 | 1029.95 | 1.00 | 0.02 |
| JY4 | 1274 | 5.02 | 1598.66 | 1547.07 | 1.00 | 0.02 |
| JY5 | 1661 | 5.21 | 2128.93 | 2128.89 | 0.99 | 0.02 |
| JY6 | 702 | 4.64 | 876.12 | 869.08 | 1.00 | 0.02 |
| JY7 | 947 | 4.42 | 1288.24 | 1270.84 | 0.99 | 0.03 |
| JY8 | 1300 | 5.43 | 1641.69 | 1604.73 | 0.99 | 0.01 |
| JY9 | 1083 | 4.78 | 1335.78 | 1314.02 | 1.00 | 0.02 |
| YS1 | 1782 | 5.68 | 2197.16 | 2261.74 | 0.99 | 0.02 |
| YS10 | 1962 | 5.75 | 2376.34 | 2345.53 | 0.99 | 0.01 |
| YS11 | 1979 | 5.62 | 2715.39 | 2557.97 | 0.96 | 0.04 |
| YS12 | 1966 | 5.70 | 2624.13 | 2506.00 | 0.97 | 0.02 |
| YS13 | 1581 | 5.34 | 1926.27 | 1899.09 | 0.99 | 0.02 |
| YS14 | 1595 | 5.79 | 1965.54 | 1964.15 | 0.99 | 0.01 |
| YS2 | 1665 | 5.75 | 2072.70 | 2139.55 | 0.99 | 0.01 |
| YS3 | 1838 | 5.50 | 2281.20 | 2383.53 | 0.99 | 0.02 |
| YS4 | 2279 | 5.92 | 2640.74 | 2548.23 | 0.99 | 0.01 |
| YS5 | 1755 | 5.69 | 2209.43 | 2257.70 | 0.99 | 0.02 |
| YS6 | 1767 | 5.51 | 2062.40 | 2061.90 | 0.99 | 0.03 |
| YS7 | 1863 | 5.20 | 2305.98 | 2374.99 | 0.99 | 0.04 |
| YS8 | 1794 | 5.87 | 2084.98 | 2069.78 | 0.99 | 0.01 |
| YS9 | 1670 | 5.54 | 2093.77 | 2071.97 | 0.99 | 0.02 |

Wild and captive roe deer were divided into two groups (A, B): captive roe deer labeled JY; wild roe deer labeledYS.

### DNA Extraction

The DNA was extracted with reference to the OMEGA-kit E.Z.N.ATM Mag-Bind Soil DNA Kit.

### DNA Extraction, PCR Amplification and Sequencing

Genomic DNA was accurately quantified using the Qubit 2.0 DNA Assay Kit to determine the amount of DNA that should be added to the PCR reaction. The first round of amplification was the V3-V4 region of the 16S rRNA gene(94°C for 3 min, followed by 5 cycles of 94°C for 30s, 45°C for 20s, 65°C for 30s, followed by 20 cycles of 94°C for 20s, 55°C for 20s, 72°C for 30s, and 72°C for 5min). The universal primers used were 341F(CCTACGGGNGGCWGCAG) and 805R(GACTACHVGGGTATCTAATCC). For the second round of amplification, we introduced Illumina bridge PCR compatible primers, and the PCR product was detected by agarose gel electrophoresis (95°C for 3 min, followed by 5 cycles of 94°C for 30s, 55°C for 20s, 72°C for 30s, and 72°C for 5min). For PCR products amplified by bacteria and archaea and normal amplified fragments, PCR products of 400 bp or more were treated with 0.6 times of magnetic beads (Agencourt AMPure XP). For fungal PCR products and PCR products with other amplified fragments of less than 400 bp, 0.8 times of magnetic beads were used for treatment. The recovered DNA was accurately quantified using the Qubit 2.0 DNA Assay Kit to facilitate sequencing in an equal amount of 1:1. When mixing in equal amounts, the amount of DNA in each sample is 10 ng, and the final concentration on the machine is 20 pmol. Finally, the machine is sequenced.

### Statistical and Bioinformatics Analyses

The raw image data files obtained by Illumina MiseqTM were converted to the original sequencing sequence (Sequenced Reads) by CASAVA base recognition(Base Calling) analysis. The Miseq sequencing sequence contains the barcode sequence, the primers and linker sequences added during sequencing. The 3’-end sequencing primer linker was removed, and the Read1 3’-end sequencing linker was TGGAATTCTCGGGTGCCAAGGAACTC. The overcap relationship between pe reads will be a sequence of “merge”. Each sample data is segmented from the merged data according to each sample barcode sequence. We removed bases with a read tail mass below 20 in each sample. We set the port of 10bp; if the average mass value in the window was less than 20, we removed the back-end base from the window. The sequence containing the N part of the reads was excised, the short sequence in the data was removed, and the length threshold was 200 bp. Then the sequences with low complexity were filtered to obtain the valid data of each sample. Usearch was removed the sequence of the non-amplified region in the pre-processed sequence, and then the sequence was sequenced for error correction, and uchime was called to identify the chimera. Subsequently, the sequence of the removed chimera is blastn-aligned with the representative sequence of the database, and the result of the alignment below the threshold was considered to be the sequence outside the target region, and the partial sequence was eliminated to obtain the final effective data.

Classification of sequences into different operation taxonomic units (OTUs) according to similarity,the similarity is greater than or equal to 97%. Using mothur for alpha diversity index analysis (Shannon, ACE, Chao1, coverage, Simpson), expressed as the means ± SD, we analyzed whether the microbiota species differences were significant by using an independent sample t-test. A P-value < 0.05 was considered statistically significant,, and P-value < 0.01 as extremely significant. Analysis of beta-diversity was performed using unweighted and weighted Unifrac,weighted unifrac and unweighted unifrac distances were used for Principal coordinate analysis(PCoA) and a one-way analysis of similarity(ANOSIM) determine differences in intestinal microflora between populations. Linear discriminant analysis Effect Size(LEfSe),The Kruskal-Wallis (KW) sum-rank test with non-parametric coefficients was used to detect the significant differences in abundance between groups. Further, unpaired Wilcoxon rank-sum test for the consistency check of the difference characteristics of the previous step in the sub-group was performed. Finally, Linear discriminant analysis(LDA) was used to estimate the effect of these differences on the differences between groups.

## ACKNOWLEDGMENTS

This work was supported by the Fundamental Research Funds for the Central Universities (2572017CA18). We thank Nathan Roberts for his valuable suggestions on retouching articles. Special thanks to all breeders in the roe deer farm (Longjia Village, Shangying Town, Shulan City).

